# Cost-effectiveness of pharmacogenetic-guided treatment: are we there yet?

**DOI:** 10.1101/065540

**Authors:** Moira Verbelen, Michael E Weale, Cathryn M Lewis

## Abstract

Pharmacogenetics (PGx) has the potential to personalize pharmaceutical treatments. Many relevant gene-drug associations have been discovered, but PGx guided treatment needs to be cost-effective as well as clinically beneficial to be incorporated into standard healthcare. Progress in this area can be assessed by reviewing economic evaluations to determine the cost-effectiveness of PGx testing versus standard treatment. We performed a review of economic evaluations for PGx associations listed in the US Food and Drug Administration (FDA) Table of Pharmacogenomic Biomarkers in Drug Labeling (http://www.fda.gov/Drugs/ScienceResearch/ResearchAreas/Pharmacogenetics/ucm083378.htm). We determined the proportion of evaluations that found PGx guided treatment to be cost-effective or dominant over the alternative strategies, and we estimated the impact on this proportion of removing the cost of genetic testing. Of the 130 PGx associations in the FDA table, 44 economic evaluations, relating to 10 drugs, were identified. Of these evaluations, 57% drew conclusions in favour of PGx testing, of which 30% were cost-effective and 27% were dominant (cost-saving). If genetic information was freely available, 75% of economic evaluations would support PGx guided treatment, of which 25% would be cost-effective and 50% would be dominant. Thus, PGx guided treatment can be a cost-effective and even cost-saving strategy. Having genetic information readily available in the clinical health record is a realistic future prospect, and would make more genetic tests economically worthwhile. However, few drugs with PGx associations have been studied and more economic evaluations are needed to underpin the uptake of genetic testing in clinical practice.

## 1 Introduction

Pharmacogenetics (PGx) studies the relationship between genetic variation and inter-individual variability in drug response in terms of efficacy and safety. Hence, PGx knowledge can be used to tailor pharmaceutical treatment to the genetic make-up of the patient. Several robust, well-replicated PGx associations exist, for example the association of *HLA–B*5701* with abacavir hypersensitivity, *HLA-B*1502* with carbamazepine induced Stevens-Johnson syndrome/toxic epidermal necrolysis (SJS/TEN), and *VKORC1* and *CYP2C9*with warfarin dosing. ^1-3^Accordingly, the US Food and Drug Administration (FDA) includes information about PGx associations in many drug labels in a wide range of therapeutic areas.^4^ These PGx drug labels cover tests that are commonly used, but also include weaker genetic associations that are reported without requiring adjustments to pharmaceutical treatment. Most drugs with mandatory genetic testing are used in oncology, but PGx tests for many therapeutic areas are already being offered by laboratories and some have become part of standard clinical practice.^5,6^

As healthcare resources are finite, it is important that the cost-effectiveness of novel PGx guided treatment strategies is assessed in addition to their clinical utility before they are widely applied. Economic evaluations, which compare costs and outcomes of at least two competing interventions, are a useful tool to inform decision making and prioritise healthcare spending. In the context of PGx testing, a pharmaco-economic study might contrast PGx guided treatment with standard treatment with the same drug, or with an alternative drug that does not require genetic testing, or with both alternatives.

Previously published literature reviews of PGx guided treatment and personalized medicine reported that the majority of PGx strategies were cost-effective or even dominant, though they noted that there was large heterogeneity in methodology between studies.^7-11^ Concerns over the quality of the early economic evaluations of PGx guided treatment have been raised, but the quality is generally considered to have improved over time.^12-15^

Our review of pharmaco-economic studies of PGx guided treatment provides an update on the literature in this rapidly evolving field (the most recent previous review covered studies up to early 2013 ^9^). Furthermore, we include a more extensive range of economic evaluations, whereas recent literature reviews were limited to cost utility analyses (CUA) only.^9,10^ We also assessed the impact of freely available genetic information on the cost-effectiveness of PGx guided treatment. We adopted a narrow definition of PGx, limiting our scope to consideration of variation in germline DNA. In contrast to tests on tumour, viral or bacterial DNA, germline DNA has the advantage that genetic variants need to be typed only once, and results remain relevant throughout a patient’s life.

## 2 Methods

### 2.1 Data sources and search strategy

The FDA Table of Pharmacogenomic Biomarkers in Drug Labeling lists FDA approved drugs that include PGx information on their drug label along with the biomarker gene (accessed on September 18, 2015).^4^ We used this table to identify drugs for which there is a genetic variant associated with the drug efficacy, safety or dosing. We excluded non-germline genetic biomarkers, for example mutations in viral or tumour DNA.

We then searched for the selected drugs and biomarkers in the National Health Service Economic Evaluations Database (NHS EED), a UK Department of Health and National Institute for Health Research funded registry of economic evaluations of health and social care interventions.^16,17^ This resource includes cost utility analyses (CUA), cost effectiveness analyses (CEA), cost benefit analyses (CBA – see below for definitions of these terms) and commentaries by the Centre for Reviews and Dissemination of the University of York. Funding of the NHS EED ceased in March 2015 and the latest database update was December 2014.

For each drug included in our study, the NHS EED was searched for economic evaluations that contain (1) the drug name and (2) the specific gene from the FDA label or the search terms *genetic*, *genotype*, *genotypic*, *pharmacogenetic* or *pharmacogenomic* in any field. We only included studies that compared a PGx guided treatment strategy with at least one alternative strategy.

We also searched PubMed to identify more recent papers (until September 2015) and any other studies missed by the NHS EED search. We searched for articles that included (1) the name of the drug and (2) the specific gene mentioned in the FDA label or the search terms *genetic*, *genotype*, *genotypic*, *pharmacogenetic* or *pharmacogenomic* in the title or abstract and (3) *Cost-Benefit Analysis* as a Medical Subject Headings (MeSH) term. In addition, the reference lists of retrieved publications were used to identify additional studies missed in our database searches.

### 2.2 Overview of economic evaluation methodology

Measuring and comparing costs and health outcomes is essential in a pharmaco-economic study. Whereas costs are naturally expressed in monetary units, the effect of a healthcare intervention can be expressed in different ways. In cost utility analyses (CUAs), health outcomes are assessed as quality-adjusted life years (QALYs), which measure the expected number of post-treatment years of life accounting for the quality of life. QALYs allow comparisons of treatment strategies across therapeutic areas and populations, but are an abstract concept (‘quality’ is hard to define) and their validity has been questioned.^18^ Cost-effectiveness analyses (CEA) evaluate the effect of an intervention in terms of a disease or treatment specific measure, for example the number of adverse events avoided, the change in score on a depression rating scale or time taken to remission. Cost-benefit analyses (CBA) quantify treatment outcome in purely monetary terms.

Furthermore, the perspective of a pharmaco-economic study determines which costs and benefits are taken into account. These can be limited to costs to the public healthcare system or private insurers, for example staff salaries, drugs and equipment costs, or may include broader costs such as productivity losses and informal care. Commonly used perspectives are the third party payer and societal perspective, but some studies take a hospital or patient perspective.

The incremental cost-effectiveness ratio (ICER) summarizes the difference in costs and health outcomes between a PGx guided strategy and standard treatment (ST):

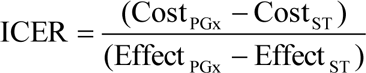

If the PGx treatment reduces costs and achieves a better outcome than the standard treatment, the PGx strategy *dominates* the standard treatment. Contrarily, if the PGx option costs more but is less effective than the standard treatment, then the PGx treatment is dominated by the standard treatment (Fig.1). When one treatment comes at a higher cost but is also more effective than the other, the ICER is compared to a willingness-to-pay threshold to determine cost-effectiveness. Generally, ICERs up to £20,000 -£30,000/QALY (or $30,000/QALY - $50,000/QALY) are considered cost-effective.^19^ As costs, health outcomes and willingness-to-pay thresholds differ between countries, or may differ according to the assumptions and perspectives adopted, economic studies evaluating the same PGx test may come to different conclusions.

**Figure 1:**
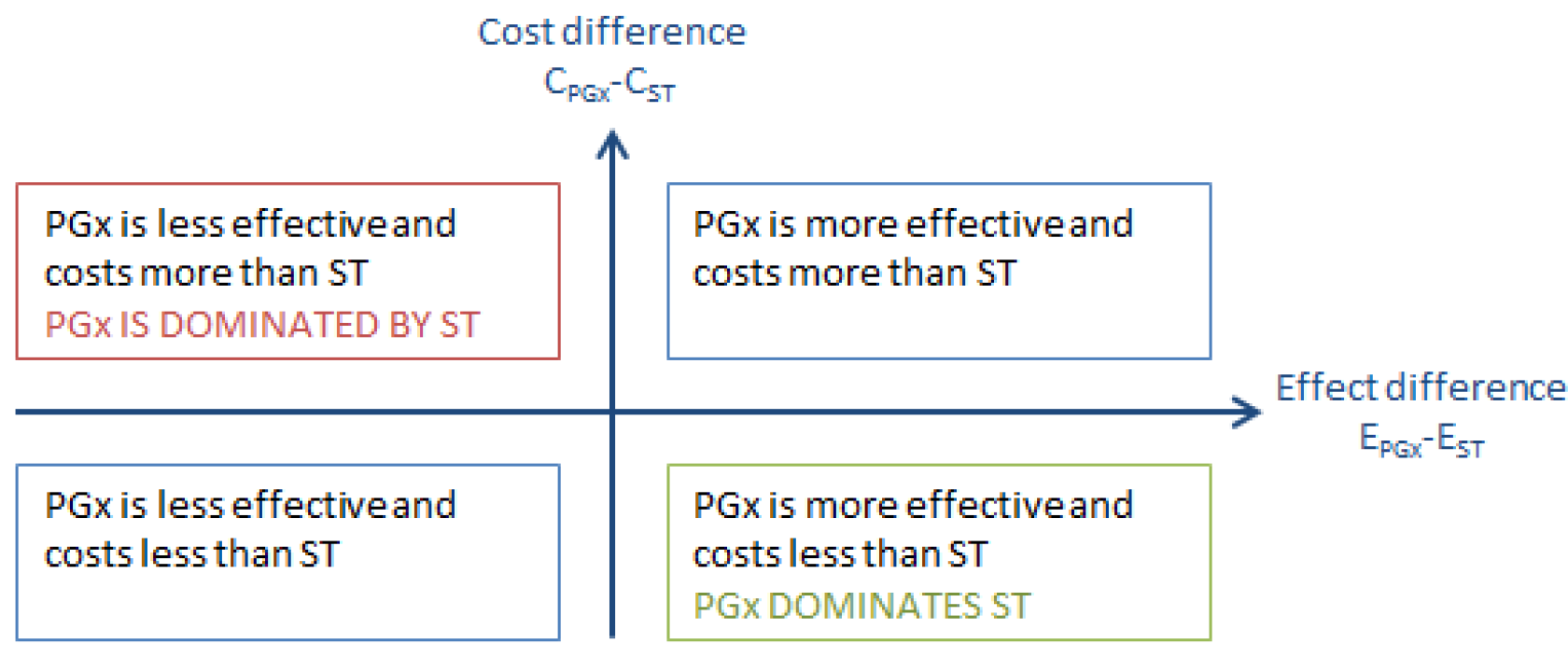
Cost-effectiveness plane of pharmaco-economic studies. PGx: pharmacogenetics guided treatment. ST: standard treatment.

### 2.3 Analyses

We extracted key parameters from the reviewed economic evaluations, including the unit of outcome, country, perspective, incremental cost-effectiveness ratio (ICER), if applicable, and the conclusion regarding the cost-effectiveness of the PGx testing strategy. A parameter of particular interest is the cost of the genetic test, as this can significantly affect the cost-effectiveness of the PGx testing strategy and may change over time. To allow comparison between studies, the price of the genetic test was corrected for inflation and converted to US dollars estimated at 2014 levels (2014 US$).

A stepwise linear regression model was fitted to test if publication year, geographic region (Asia, Oceania, US and Canada or EU) or perspective (healthcare, society or other) had an influence on the price of genetic testing. A stepwise logistic regression model was also used to investigate whether publication year, geographic region, perspective, cost of genetic test, genetic variant (*HLA*, *TPMT* or other), or outcome (QALY or other) was associated with the PGx testing strategy being cost-effective. Statistical analysis was performed in R (version 3.1.2).

We estimated the impact of freely available genetic information on the conclusions regarding the cost-effectiveness of PGx informed strategies. The ICER under assumption of free genetic testing was calculated by adjusting the cost of the PGx guided treatment by the cost of the test

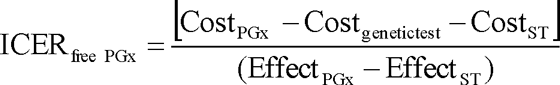

When insufficient details were provided to estimate the ICER _free PGx_ , it was assumed that free genetic testing could not worsen the conclusion regarding PGx guided treatment. For example, when a study found the PGx strategy to be cost-effective, we assumed that PGx guided treatment with free genetic testing would also be at least as cost-effective.

## 3 Results

### 3.1 Description of studies

The FDA Table of Pharmacogenomic Biomarkers in Drug Labeling listed 137 distinct drugs, of which 68 met our inclusion criteria (Fig. 2). These drugs were from diverse clinical specialties, including cancer (11 drugs), infectious diseases (10 drugs), psychiatry (9 drugs) and neurology (8 drugs) (Table 1). Our literature search yielded economic evaluations for only 10 of these 68 drugs (14.7%) (Table 2). All publications related to a single drug, except for one study investigating a PGx testing strategy for carbamazepine and phenytoin treatment, which assumed both drugs to be interchangeable in terms of costs, efficacy and safety.^20^ To avoid duplication of studies in our review, this publication was counted as a carbamazepine study (there were no other publications on phenytoin).

**Figure 2:**
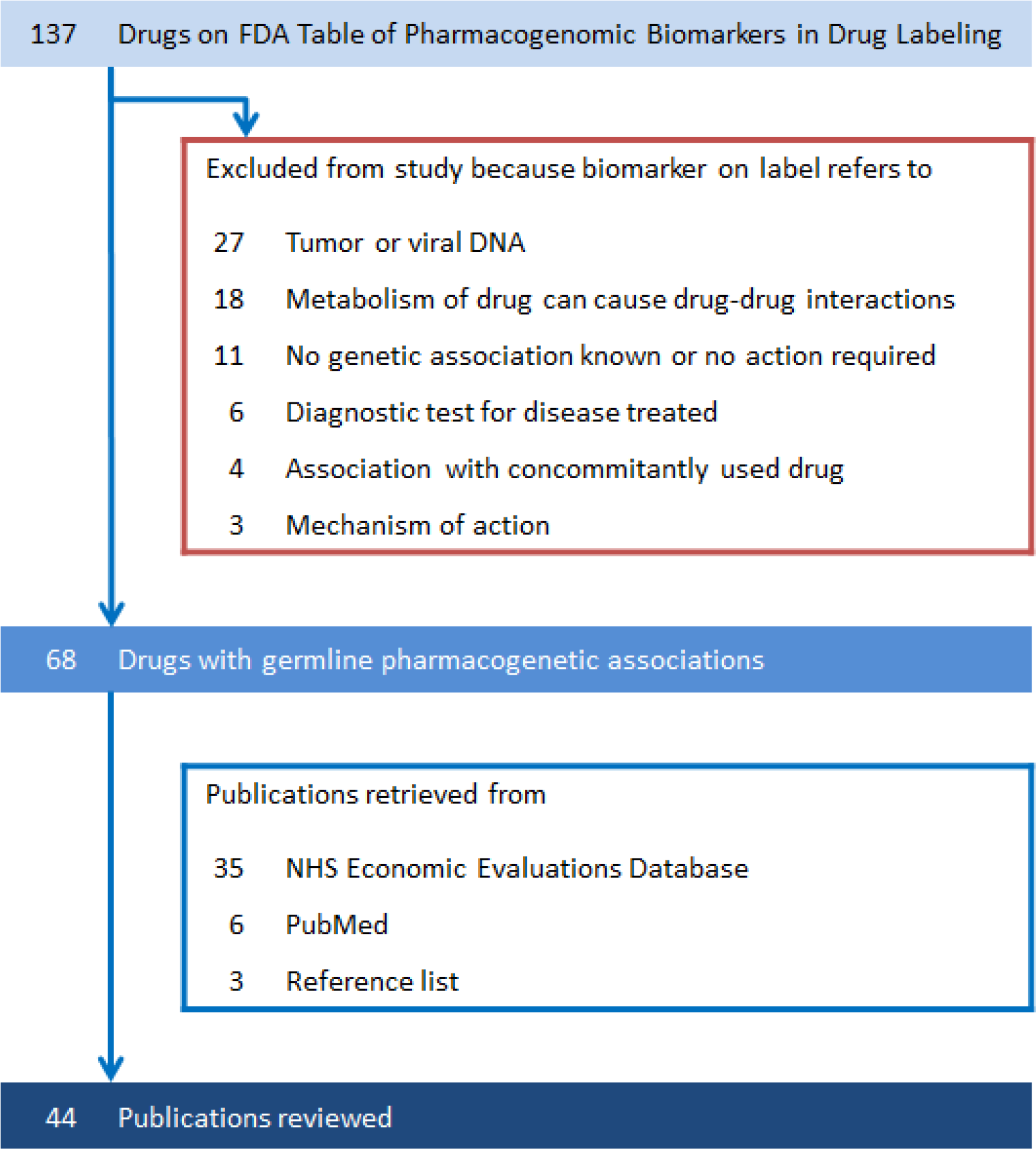
Number of drugs and publications included in literature review.

**Table 1:**
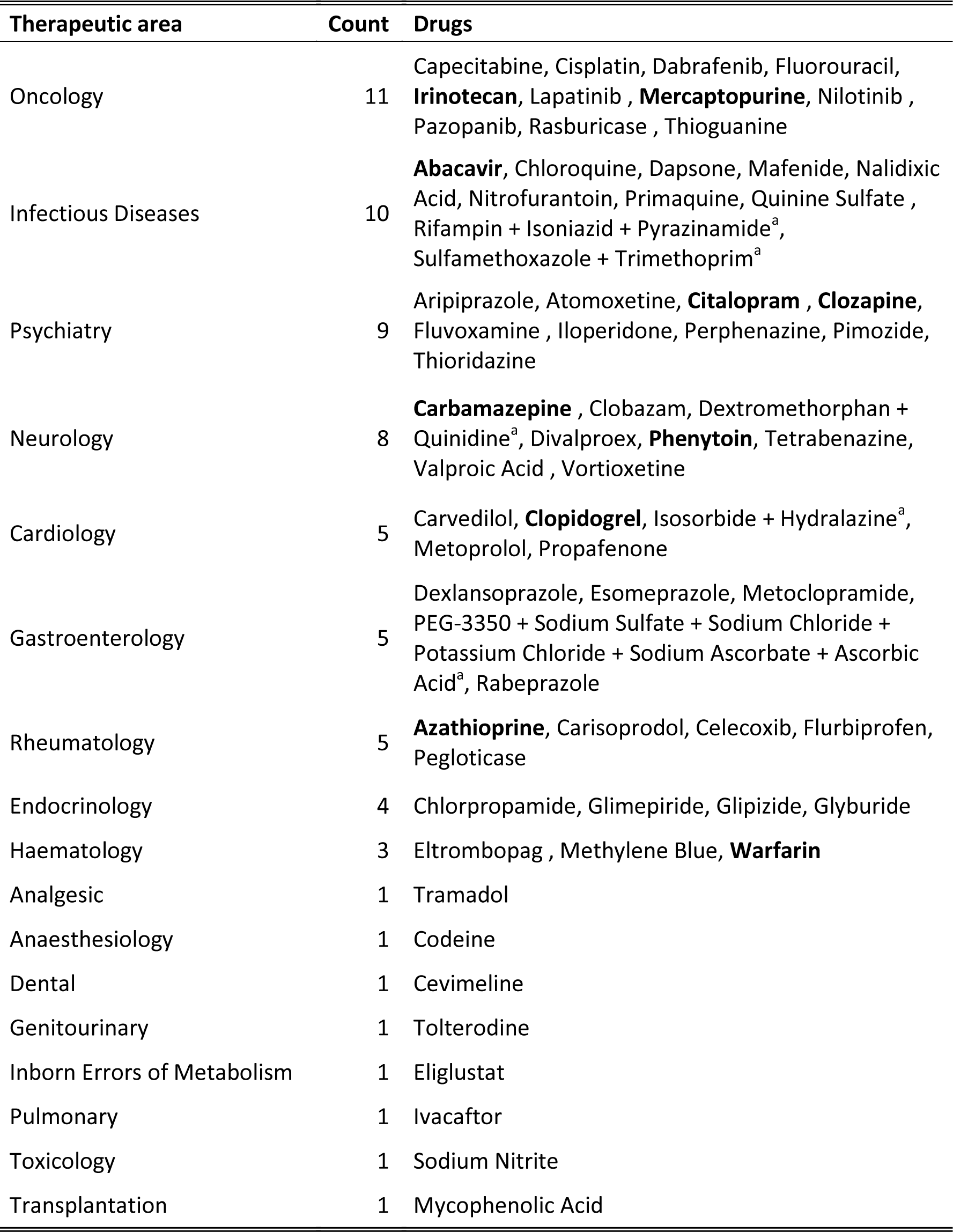
Drugs from FDA Table of Pharmacogenomic Biomarkers in Drug Labeling included in literature review. Drugs in bold had economic evaluations available.

| Therapeutic area | Count | Drugs |
| --- | --- | --- |
| Oncology | 11 | Capecitabine, Cisplatin, Dabrafenib, Fluorouracil, **Irinotecan**, Lapatinib , **Mercaptopurine**, Nilotinib , Pazopanib, Rasburicase , Thioguanine |
| Infectious Diseases | 10 | **Abacavir**, Chloroquine, Dapsone, Mafenide, Nalidixic Acid, Nitrofurantoin, Primaquine, Quinine Sulfate , Rifampin + Isoniazid + Pyrazinamide<sup>a</sup>, Sulfamethoxazole + Trimethoprim<sup>a</sup> |
| Psychiatry | 9 | Aripiprazole, Atomoxetine, **Citalopram** , **Clozapine**, Fluvoxamine , Iloperidone, Perphenazine, Pimozide, Thioridazine |
| Neurology | 8 | **Carbamazepine** , Clobazam, Dextromethorphan + Quinidine<sup>a</sup>, Divalproex, **Phenytoin**, Tetrabenazine, Valproic Acid , Vortioxetine |
| Cardiology | 5 | Carvedilol, **Clopidogrel**, Isosorbide + Hydralazine<sup>a</sup>, Metoprolol, Propafenone |
| Gastroenterology | 5 | Dexlansoprazole, Esomeprazole, Metoclopramide, PEG-3350 + Sodium Sulfate + Sodium Chloride + Potassium Chloride + Sodium Ascorbate + Ascorbic Acid<sup>a</sup>, Rabeprazole |
| Rheumatology | 5 | **Azathioprine**, Carisoprodol, Celecoxib, Flurbiprofen, Pegloticase |
| Endocrinology | 4 | Chlorpropamide, Glimepiride, Glipizide, Glyburide |
| Haematology | 3 | Eltrombopag , Methylene Blue, **Warfarin** |
| Analgesic | 1 | Tramadol |
| Anaesthesiology | 1 | Codeine |
| Dental | 1 | Cevimeline |
| Genitourinary | 1 | Tolterodine |
| Inborn Errors of Metabolism | 1 | Eliglustat |
| Pulmonary | 1 | Ivacaftor |
| Toxicology | 1 | Sodium Nitrite |
| Transplantation | 1 | Mycophenolic Acid |

<sup>a</sup> Multiple drugs on a single FDA label.

**Table 2:** Drugs for which a PGx guided strategy was studied in economic evaluation(s).

| Drug | Therapeutic area | Gene | Notes (based on PharmGKB.org<sup>32</sup>) | Number of reviewed publications |
| --- | --- | --- | --- | --- |
| Abacavir | HIV | *HLA-B* | Abacavir is contraindicated for *HLA-B\*5701* carriers as they are at high risk of hypersensitivity reaction. | 5 (ref. 33-37) |
| Azathioprine | Rheumatology | *TPMT* | Carriers of 1 nonfunctional *TPMT* allele may require reduced azathioprine dose. Carriers of 2 nonfunctional *TPMT* alleles are at high risk of myelotoxicity and alternative treatment should be considered. | 9 (ref. 38-46) |
| Carbamazepine | Neurology | *HLA-B, HLA-A* | Carbamazepine is contraindicated for *HLA-B\*1502* carriers as they are at high risk Stevens-Johnson Syndrome/toxic epidermal necrolysis. *HLA-A\*3101* has also been associated with hypersensitivity reactions. | 4 (ref. 20,47-49) |
| Citalopram | Psychiatry | *CYP2C19, 5-HTTLPR<sup>a</sup>, HTR2A<sup>a</sup>* | *CYP2C19* poor metabolizers require reduced citalopram starting dose. Polymorphisms in 5-HTTLPR and HTR2A are associated with citalopram response.<sup>50,51</sup> | 3 (ref. 52-54) |
| Clopidogrel | Cardiology | *CYP2C19* | *CYP2C19* poor metabolizers have reduced response to clopidogrel and alternative treatment should be considered. | 6 (ref. 24,25,55-58) |
| Clozapine | Psychiatry | *CYP2D6, H2<sup>a</sup>, 5-HTT<sup>a</sup>, 5-HT<sub>2A</sub><sup>a</sup>, 5-HT<sub>2C</sub><sup>a</sup>* | *CYP2D6* poor metabolizers may require reduced clozapine dose. Six polymorphisms in H2, 5-HTT, 5-HT2A and 5-HT2C are associated with clozapine response.<sup>59</sup> | 1 (ref. 60) |
| Irinotecan | Oncology | *UGT1A1* | Patients homozygous for the *UGT1A1\*28* allele are at higher risk of neutropenia and should receive a reduced starting dose of irinotecan. | 3 (ref. 61-63) |
| Mercaptopurine | Oncology | *TPMT* | Carriers of 1 nonfunctional *TPMT* allele may require reduced mercaptopurine dose. Carriers of 2 nonfunctional *TPMT* alleles are at high risk of myelotoxicity and alternative treatment should be considered. | 1 (ref. 64) |
| Warfarin | Cardiology | *CYP2C9, VKORC1* | Genetic variation in *VKORC1* and *CYP2C9* explain 40% variance in warfarin dose. Genetic and clinical information can be used to determine starting dose. | 12 (ref. 21-23,65-73) |

We retrieved 44 economic evaluations that investigated the cost-effectiveness of a PGx informed strategy (Table 2). Full details of the reviewed studies and extracted information are given in Supplementary Table 1. The earliest study included was published in 2000 and over 70% of studies were published in 2009 or later. Most publications were CUAs (30 studies, 68%) or CEAs (12 studies, 27%), with only two CBAs (5%). A healthcare system perspective was adopted in 18 studies (41%), a societal perspective in 10 papers (23%), a third-party payer perspective in 5 studies (11%) and 11 papers (25%) did not state a clear perspective. Twenty studies (45%) were conducted in North America, 11 (25%) in Europe, 6 (14%) in Asia and 3 (7%) in Oceania; 4 studies (9%) did not specify a country. Warfarin had the most economic evaluations (12 studies), followed by azathioprine (9 studies); clozapine and mercaptopurine had only 1 economic evaluation each (Table 2).

### 3.2 Cost-effectiveness of PGx informed treatment

We assessed the overall conclusions regarding cost-effectiveness of each PGx study. Over half of the 44 economic evaluations took a favourable view of the PGx guided strategy: in 12 studies (27%) it was dominant (cost-saving) and in 13 studies (30%) it was cost-effective. Eleven publications (25%) found PGx testing not cost-effective and 8 studies (18%) were undetermined (Fig. 3a). The majority of economic evaluations concluded in favour of PGx testing for azathioprine (7 out of 9 studies), clopidogrel (4 out of 6 studies), abacavir (4 out of 5 studies), carbamazepine (3 out of 4 studies), irinotecan (3 out of 3 studies) and clozapine (1 study) (Fig. 3b). Although Warfarin had the highest number of economic studies, they reached diverging conclusions: 3 studies found PGx guided dosing cost-effective, 4 studies were undetermined and 5 studies concluded it was not cost-effective. No studies found unequivocally that PGx guided citalopram (3 studies) or mercaptopurine (1 study) treatment was cost-effective.

**Figure 3:**
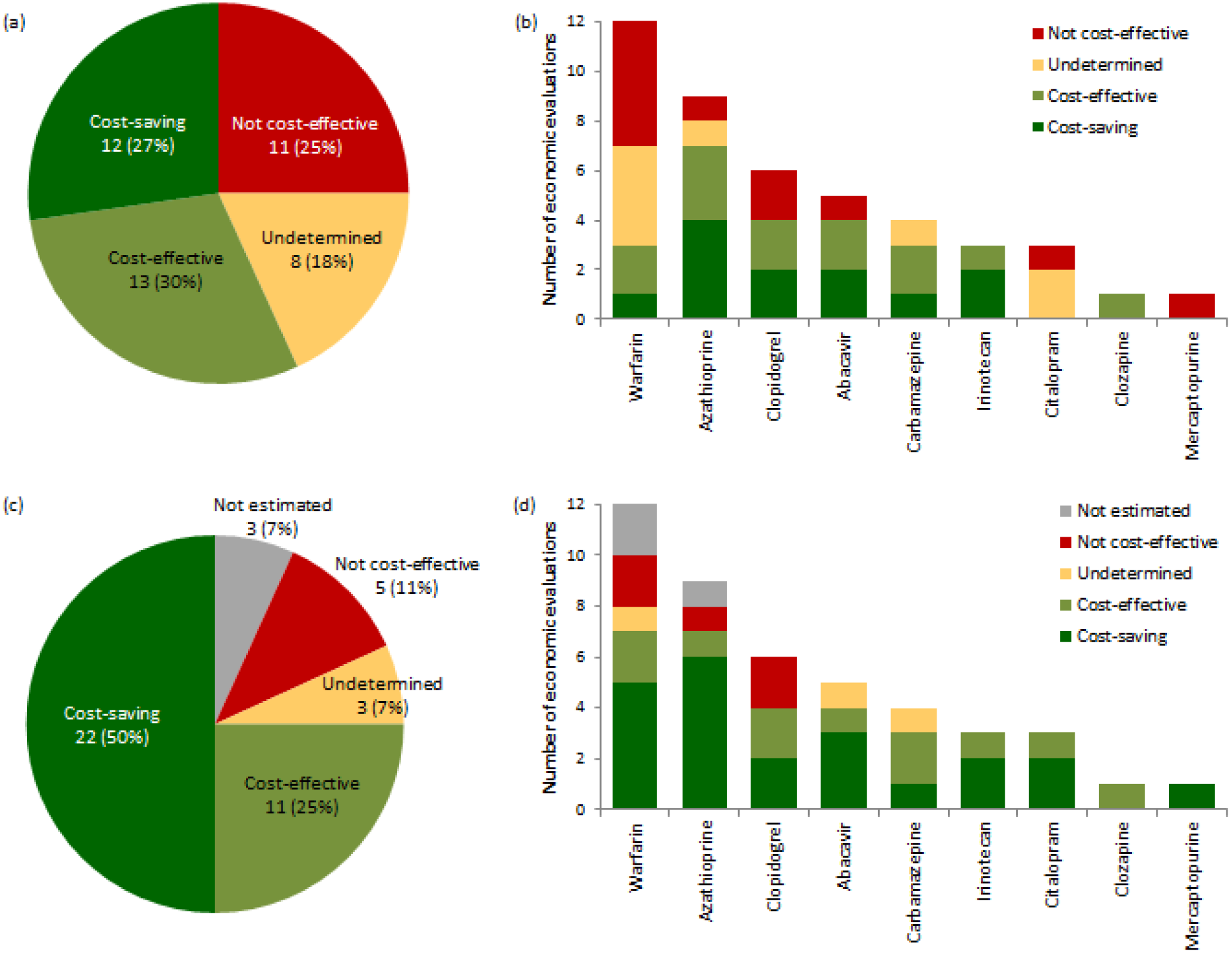
Conclusions of reviewed economic evaluations regarding cost-effectiveness of PGx testing strategy (a) overall and (b) by drug, and estimated conclusions in scenario of no extra cost for genetic information (c) overall and (d) by drug.

We assessed the effect of study characteristics on the probability of concluding in favour of the PGx strategy. A logistic regression model detected that CUAs (studies using QALYs as outcome measure) were less likely than CEAs and CBAs to find the genetic testing strategy cost-effective (odds ratio = 0.13, p-value < 0.05). However, there is no clear explanation for this and it may be a spurious result due to the relatively small sample size of 44 economic evaluations.

### 3.3 Effect of cost of genetic test on cost-effectiveness of PGx informed treatment

The cost of genetic testing is an important parameter of economic evaluations of PGx interventions. After correcting for inflation and converting to 2014 US$, the cost of genetic testing quoted by the reviewed studies ranged between US$33 and US$710 with a median value of US$175. The price of genetic tests decreased slightly over time (not statistically significant) and this trend was more pronounced since 2009, the period when most economic evaluations were published (p-value < 0.05) (Fig. 4). Prices were on average higher in the US and Canada than other regions of the world (mean US and Canada: US$363.65, mean other regions: US$131.80, p-value < 0.05). We noted a wide variability in prices of tests for the same drug. For example, the lowest price quoted for warfarin PGx testing was US$36 in a 2014 UK based study^21^, while US$600 and US$657 were used in a 2013 Canadian and 2009 US study, respectively.^22,23^ The prices for clopidogrel PGx testing also varied considerably: from US$45 (2013 Australian study) to US$ 543 (2013 US study).^24,25^

**Figure 4:**
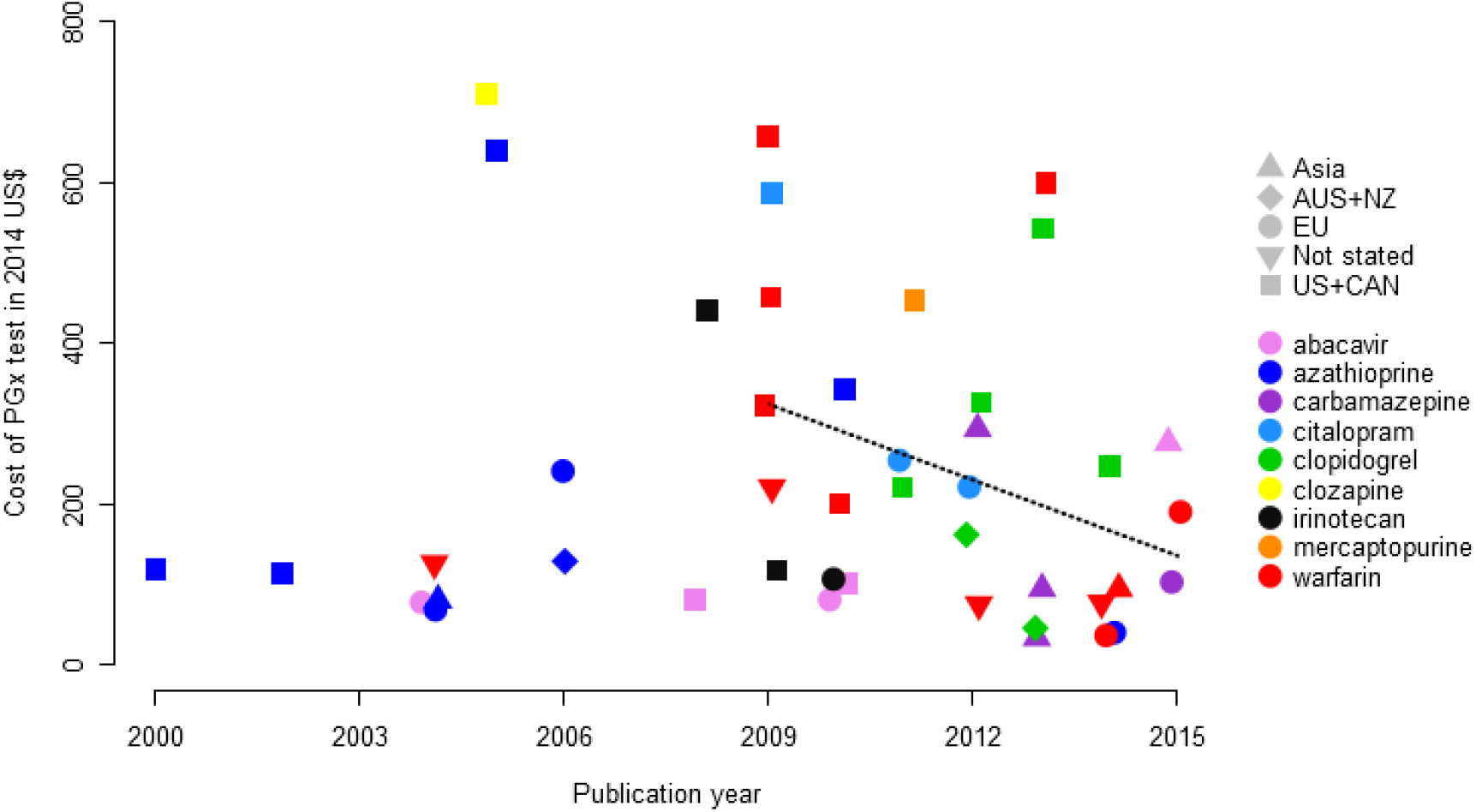
Cost of PGx test as reported in the reviewed economic evaluations over time, with fitted regression since 2009 (dotted line).

Given the decreasing costs of genetic testing and its increasing availability, we looked ahead to a possible future where genotype information might be readily available, at negligible cost, for all patients as part of their electronic health record. Thirty-three economic evaluations (75%) would support PGx guided treatment under this scenario, with 11 studies (25%) finding it cost-effective and 22 studies (50%) considering it dominant and cost-saving (Fig. 3c). Five studies (11%) would still conclude that PGx testing was not cost-effective, while 3 studies (7%) would be inconclusive. A separate set of 3 studies had to be excluded, because the impact of free genetic testing could not be estimated. Evaluating the effect of freely available genetic information by drug, we found that PGx informed treatment with citalopram and mercaptopurine would be considered cost-effective in all cases, whereas none of the reviewed studies previously concluded in favour of PGx testing (Fig. 3d). For the 12 economic evaluations of warfarin, the number of cost-effective studies would increase from 3 to 7 with freely available genetic testing.

## 4 Discussion

We have assessed published economic evaluations comparing the cost effectiveness of pharmacogenetic guided treatment to standard treatment for drugs listed in the FDA’s Table of Pharmacogenomic Biomarkers in Drug Labeling. The economic evaluations were drawn from the NHS EED database, which includes economic evaluations up to 31^st^ December 2014. An alternative source of economic studies would be the Cost-Effectiveness Registry (CEA Registry) maintained by the Tufts Medical Centre. We opted to use the more comprehensive NHS EED since the CEA Registry is limited to CUAs (measuring health outcomes in QALYs), which would have reduced the number of evaluations available for assessment. Moreover, CEA Registry was not updated beyond 2014 and it only provides advanced database searches for subscribers and contributors.^17^ A third resource, the Health Economic Evaluations Database (HEED) curated by John Wiley & Sons, was discontinued in 2014.^26^ As economic evaluations provide evidence for the introduction of PGx testing into clinical practice, we argue that an up-to-date, accessible database would be an important and valuable resource for both health-economic and PGx research.

Few of the FDA-listed drugs have been the subject of published economic evaluations assessing the economic aspects of PGx testing. This was previously also noted by Phillips et al.^10^, who found that only 13% of drugs on the FDA table and 27% of available genetic tests were accompanied by economic studies. Since healthcare budgets are limited, it is increasingly the case that clinical utility alone is not sufficient to recommend application of a PGx test in clinical practice. Thus, economic feasibility must also be assessed, and found to be favourable, before we can expect genetic tests to be widely adopted. We call for more pharmaco-economic studies in this field, and these should also be regularly updated to respond to the changing landscape of healthcare and, above all, genetic testing costs.

A limitation of our study is that the economic evaluations reviewed may not be representative of all PGx tests. The economic aspects of PGx guided treatment may not have been studied for drugs where testing is obviously necessary, for example because it prevents life-threatening adverse events or because the drug is only effective in carriers of certain genetic variants. Another possibility is that economic studies focus on PGx tests that are already applied in clinical practice and for which there is an apparent interest. There may also be a publication bias in that studies which find genetic testing to be not cost-effective may be less likely to be published. Some assumptions of the economic evaluations may have changed since publication; in particular, prices for PGx tests quoted by the reviewed studies are likely to have decreased. These constraints should be considered in the interpretation of findings of the cost-effectiveness of PGx testing based on the available literature.

The PGx dosing algorithm for warfarin is often presented as the poster child for the achievements of PGx, because the drug is widely prescribed and implementation of this single nucleotide polymorphism (SNP)-based test could have a major impact on healthcare. However, only one quarter of studies considered genetic guided dosing for warfarin to be cost-effective, and the clinical advantage of genetic guided dosing over standard dosing appears to be small or even non-existent.^27^ Although freely available genetic testing would improve the cost-effectiveness of genotype-guided warfarin dosing, other drugs such as abacavir, where genetic testing for *HLA-B*5701* is required by the FDA, might make more convincing PGx success stories.^28^

Our study assessed the characteristics of tests in the reviewed evaluations. We noted that quoted prices for genetic tests in the US and Canada were higher than in other countries, although there was also a large between-study variability within these countries. The drug tested and study perspective were not significantly associated with price. Genetic test costs may depend on the method used to determine genetic variants (e.g. polymerase chain reaction or measuring enzyme activity), but the reviewed studies did not provide sufficient detail to investigate the impact of this parameter on price. A downward trend in prices for genetic testing is apparent in recent years, and this may continue as new genetic technologies become more accessible and lead to further price reductions.

We show that the cost of genetic testing is an important factor in determining the cost-effectiveness of a PGx-guided treatment strategy. If there was no cost attached to genetic testing, the number of economic evaluations that found the PGx strategy cost-effective increased greatly, such that half of the reviewed studies considered it dominant over the alternative and 75% considered it cost-effective. Freely available genetic testing might be achievable in future as genomic prices fall and the perceived or actual value of genetic information increases. Once genetic tests become a mainstream clinical service, economies of scale will decrease the price of testing still further. For example, the direct to consumer testing company 23andMe offers a genome-wide genotyping service for £125 (UK, March 2016 price), which includes SNP-based testing for 5 of the 10 drugs covered in this review.^29^ Similarly, the cost of whole genome sequencing has fallen every year and is now in the region of US$1,000.^30^ Having genetic information in the electronic health record would allow PGx information to be queried for any new prescription or review of dosage. A genetic test would need to be performed only once and this information, safely secured and immediately accessible, could guide treatment throughout the patient’s life.

Even so, PGx guided treatment will not be cost-effective in all situations. Even under the favourable assumption of freely available genetic testing, it could be more expensive than the alternative strategy. Although this may be counter-intuitive, genetic testing costs may only be a small part of the costs attached to PGx informed treatment, so eliminating the cost of the test itself may not have a large impact. Increased costs may arise where the alternative drug for test-positive patients is much more expensive, and this is exacerbated if the test has a high proportion of false positive results. For example, *CYP2C19* poor metabolizers are prescribed the more expensive ticagrelor in place of clopidogrel (which is metabolized into its active form by *CYP2C19*), and the drug cost may out-weigh the costs incurred by the lack of efficacy of clopidogrel in the poor metabolizers.^24^Thus even if genetic information is freely accessible, economic evaluations of PGx testing are still relevant and necessary.

In general, the economic evaluation studies reviewed here showed that pharmacogenetics has a positive impact on healthcare quality and costs. Over half of reviewed studies concluded that the PGx informed treatment strategy is more cost-effective than the alternatives considered under present-day economics. Only one in four economic evaluations found the genetic testing option unequivocally not cost-effective. This encouraging finding, with an even bigger projected benefit under low-cost genetic typing, suggests that PGx testing has the potential to be a cost-effective or even cost-saving intervention. It therefore seems likely that PGx testing will become a core clinical service, particularly as projects such as the 100,000 Genomes Project pushes genomics to become part of health care infrastructure and as electronic health records become increasingly effective.^31^

Supplementary information is available at the European Journal of Human Genetics website.

## Acknowledgements

This study was funded by an industrial CASE studentship to Moira Verbelen from the Medical Research Council with Eli Lilly and Company Ltd. This paper represents independent research part-funded by the National Institute for Health Research (NIHR) Biomedical Research Centre at South London and Maudsley NHS Foundation Trust and King’s College London. The views expressed are those of the authors and not necessarily those of the NHS, the NIHR or the Department of Health.

